# HAPI Cells are SIM-A9-related Mouse Microglial Cells Useful for *In Vitro* Modeling of Microglial Immunometabolism

**DOI:** 10.64898/2026.02.11.705385

**Authors:** Ryan P. Mayers, Sausan M. Jaber, Nicolas Verhoeven, Ayodele Jaiyesimi, Brian M. Polster

## Abstract

Highly aggressively proliferating immortalized (HAPI) cells were initially described as a spontaneously immortalized rat cell line isolated from a mixed neonatal rat glial population. It was demonstrated that HAPI cells are phagocytic, stain for macrophage-/microglia-specific markers like CD11b and GLUT5, and exhibit lipopolysaccharide (LPS)-induced nitric oxide (NO) and tumor necrosis factor-alpha (TNF-α) release. These characteristics led to their widespread use as a rat microglial cell line. Here, we report that HAPI cells are mouse cells, not rat cells, but further establish that they have a microglia-like identity and properties useful for *in vitro* modeling. Cell line authentication by short tandem repeat (STR) profiling, a method that detects identifying DNA signatures, indicates that HAPI cells are a 100% match for SIM-A9 cells, a mouse microglial cell line reported to be spontaneously immortalized from primary cell culture. We find that both HAPI cells and SIM-A9 cells express the microglia-selective gene *Tmem119*, as well as the microglia-/macrophage-selective marker *Cx3cr1*, supporting a microglial origin. Like primary rodent microglia or macrophages, HAPI cells respond to combined stimulation with LPS and the Type II interferon, interferon-gamma (IFN-γ), with a pro-inflammatory morphology, NO production, NO-dependent suppression of mitochondrial oxygen consumption, and increased extracellular acidification (an indicator of glycolysis). The Type I interferon, interferon-alpha (IFN-α), also reduces mitochondrial oxygen consumption when administered alone or in combination with LPS. Overall, results indicate that HAPI cells are SIM-A9-related mouse cells of microglial origin and support their continued use to study microglial behavior *in vitro*, including immunometabolism.

## Introduction

Highly aggressive proliferating immortalized (HAPI) cells were first reported in 2001, when they were isolated from a microglia-enriched culture of mixed glial cells prepared from a three-day old rat brain (Cheepsunthorn et al., 2001). The initial report described HAPI cells as spontaneously immortalized rat microglia based on their purported rat brain origin and three characteristics shared with *in vivo* microglia. First, immunocytochemistry was used to demonstrate that HAPI cells express markers consistent with microglial or macrophage identity, such as glucose transporter type 5 (GLUT5), but not those associated with astrocytes, such as glial fibrillary acidic protein (GFAP). Second, it was shown that HAPI cells are capable of phagocytosis. Third, like rat microglia, HAPI cells responded to stimulation with the toll-like receptor 4 (TLR4) agonist lipopolysaccharide (LPS) with increased inducible nitric oxide synthase (iNOS) and tumor necrosis factor-alpha (TNF-α) expression, as well as with subsequent nitric oxide (NO) and TNF-α release (Cheepsunthorn et al., 2001).

It is difficult to obtain primary rodent microglia in sufficient numbers for many cell biology and biochemistry-based approaches, and the intractability of primary cells to genetic manipulation is another limitation. With their high proliferation rate, primary-cell-like pro-inflammatory responses, and relatively few microglial cell lines available as alternatives, the HAPI cell model has proven useful for microglial research. The findings that HAPI cells are capable of phagocytosis (Mairuae et al., 2011) and produce NO and other pro-inflammatory molecules in response to LPS and the type II interferon, interferon-gamma (IFN-γ), have been independently replicated many times, supporting the use of HAPI cells for studying microglial immune responses (Chamniansawat and Chongthammakun, 2015; Chuang et al., 2013; Dallas et al., 2013; Hara et al., 2014; Hara et al., 2024; Hara et al., 2011; Horvath et al., 2008; Jantaratnotai et al., 2022; Jantaratnotai et al., 2006; Jantaratnotai et al., 2013; J. Li et al., 2022; Lo et al., 2008; McFarland et al., 2018; Pawate et al., 2004; Sheng et al., 2011; Tocharus et al., 2012).

Since the HAPI cell line’s initial description, the field of immunometabolism has emerged, which studies the relationship between altered cellular metabolism and immune behavior. Primary microglia treated with the pro-inflammatory stimulus LPS, alone or in combination with IFN-γ, exhibit suppressed mitochondrial respiration and elevated glycolysis (Jaber et al., 2018; Orihuela et al., 2016), however, the biochemical mechanisms underlying these bioenergetic changes are incompletely understood. Because of the aforementioned limitations of primary microglia, we initially sought to establish whether HAPI cells exhibit primary microglia-like immunometabolic changes and could be used to investigate these mechanisms using a genetic approach. However, while conducting CRISPR gene editing experiments, we performed next-generation sequencing of multiple genes and the results suggested that HAPI cells have a mouse, rather than rat, origin. We subsequently discovered that HAPI cells are listed on the International Cell Line Authentication Committee’s Register of Misidentified Cell Lines as mouse, not rat, cells based on personal communication with their former suppliers, MilliporeSigma (Burlington, MA). Although HAPI cells first appeared in version 11 of this database, which was released on June 8, 2021, we have not identified any peer-reviewed publications referring to HAPI cells as mouse cells, and recent publications continue to refer to HAPI cells as rat microglia (Albalawi et al., 2024; Cao et al., 2023; Gao et al., 2024; Guo et al., 2025; Y. Li et al., 2023; Peng et al., 2024; Shen et al., 2023; Shukitt-Hale et al., 2025; Wang et al., 2024; Wei et al., 2024; Zhu et al., 2023).

A “HAPI microglia*” PubMed search now retrieves over 100 results that include studies relevant to a range of neurodegenerative diseases and other neurological conditions. Given this significant body of work, the goals of our study became: 1) to authenticate HAPI cells as a microglia-like *mouse* cell line using the gold-standard method of short tandem repeat (STR) profiling and expression studies of the newer, widely used microglia-selective marker, transmembrane protein 119 (TMEM119), and 2) to help re-establish and extend the utility of HAPI cells for *in vitro* neuroinflammation studies by measuring the morphological and bioenergetic changes induced by disease-relevant pro-inflammatory stimuli, i.e., the TLR4 agonist LPS, the Type I interferon, IFN-α, and the Type II interferon, IFN-γ.

We found that the HAPI DNA signature revealed by STR profiling was surprisingly identical to that of mouse SIM-A9 microglial cells and other approaches provided further support for a microglial origin.

## Materials and Methods

### Materials

Universal™ Type I interferon, a hybrid human interferon-alpha (IFN-α) protein exhibiting bioactivity across multiple species including mice, was from PBL Assay Science (Piscataway, NJ; catalogue # (Cat#) 11200). Mouse-origin recombinant IFN-γ protein was from Sino Biological (Paoli, PA; Cat# 50709-MNAH). Rat-origin recombinant IFN-γ protein was from Thermo Fisher Scientific (Waltham, MA; Cat# 400-20). All other reagents were purchased from Sigma-Aldrich (St. Louis, MO) unless otherwise noted.

### Animals for Tissue Isolation and Primary Cell Culture

Adult male C57BL6/J mice were kept on a 12 hour light/dark cycle, fed *ad libitum*, and housed according to standard animal care protocols. All procedures were consistent with the National Institutes of Health *Guide for the Care and Use of Laboratory Animals* and approved by the University of Maryland Baltimore Institutional Animal Care and Use Committee.

### Cell Culture

HAPI cells were generously provided by Dr. James Connor (Pennsylvania State University; Hershey, PA) and Neuro2a cells were kindly provided by Dr. Philip Iffland (University of Maryland School of Medicine; Baltimore, MD). SIM-A9 cells (Cat# CRL-3265) and LADMAC cells (Cat# CRL-2420) were purchased from the American Type Culture Collection (ATCC; Manassas, VA).

HAPI and SIM-A9 cells were cultured in Dulbecco’s Modified Eagle Medium (DMEM) supplemented with 10% heat-inactivated fetal bovine serum (FBS) and 100 IU/ml penicillin plus 100 μg/ml streptomycin (P/S), whereas Neuro2a cells and LADMAC cells were cultured in Eagle’s Minimum Essential Medium supplemented with 10% heat-inactivated FBS and P/S. Only cell line passages under 30 were utilized. Bone-marrow derived macrophages (BMDMs) were isolated from male C57BL/6 mice as described (Haag and Murthy, 2021) and cultured for eight days in DMEM supplemented with 10% heat-inactivated FBS, P/S, and 20% DMEM conditioned from LADMAC cells. A full media change was performed every other day. All cells were maintained in 95% air and 5% CO2 at 37°C.

HAPI cells were seeded in V7 XF24 plates (Agilent Technologies, Santa Clara, CA) at 7,500-15,000 cells per well (depending on the individual passage’s recent growth rate) for Seahorse extracellular flux experiments and allowed to recover from plating overnight prior to treatment.

### Short Tandem Repeat (STR) Profiling

A HAPI cell pellet was submitted to the ATCC for DNA extraction followed by STR profiling. Eighteen mouse STR loci were analyzed by ATCC and two markers (Human D8 and D4) were used to screen for the presence of human or African green monkey species. Samples, including appropriate positive and negative controls, were processed using an ABI Prism® 3500xl Genetic Analyzer and data were analyzed using GeneMapper® ID-X v1.2 software (Applied Biosystems; Waltham, MA). Comparison to entries in the ATCC Mouse STR Database were conducted by ATCC according to the Tanabe algorithm (Tanabe et al., 1999).

### Immunocytochemistry

HAPI cells were seeded on 1.5H glass coverslips at 75,000 cells per well in 24-well plates and cultured overnight. The cells were then fixed on the following day in 4% paraformaldehyde in phosphate-buffer saline (PBS) for 20 minutes. The fixed cells were washed three times with PBS, permeabilized with 0.15% Triton-X-100 in PBS for 20 minutes, and blocked with 7.5% bovine serum albumin (BSA) in PBS for 45 minutes. Primary immunostaining was performed with a 1:200 dilution of mouse anti-TMEM119 antibody (Thermo Fisher Scientific, Cat# MA5-35043, RRID:AB_2848948) for 90 minutes in PBS containing 7.5% BSA and 0.15% Triton-X-100. Cells were then washed twice in PBS, before 60 minutes of incubation with an Alexa Fluor 488 goat anti-mouse IgG secondary antibody (Thermo Fisher Scientific, Cat# A11029, RRID:AB_2534088) at a 1:1000 dilution. Cells were then washed twice more in PBS before mounting coverslips and allowing to cure overnight in ProLong™ Diamond Antifade Mountant with DAPI (Thermo Fisher, Cat# P36971). Imaging was performed on a Nikon W1 spinning disk confocal (Nikon Instruments Inc., Melville, NY) mounted on a Nikon Ti2 inverted microscope at the University of Maryland, Baltimore Confocal Microscopy Facility. All steps were performed at room temperature.

### RNA Extraction and Polymerase Chain Reaction (PCR)

The cell lines were plated at 300,000 cells/well in 6-well plates and BMDMs were plated in three 10 cm^2^ dishes for RNA extraction; three wells or dishes were combined for each extraction. RNA from HAPI cells, SIM-A9 cells, mouse cortex, mouse BMDMs, or Neuro2a cells was extracted and isolated using an RNAeasy Minikit (Cat# 74104) from Qiagen (Germantown, MD) according to the manufacturer’s protocol. cDNA amplification was performed using a High-Capacity cDNA Reverse Transcription Kit (Thermo Fisher Scientific, Cat# 4368814) according to the manufacturer’s protocol, with 300 ng of input RNA for each sample. Primers against *Tmem119* (expected product length: 119 base pairs (bp)) and *Cx3cr1* (expected product length: 109 bp) were purchased from Integrated DNA Technologies (Coralville, IA). The *Tmem119* forward and reverse primers were ATGGGGACAATGACAGCTCTTC and GGGAGTGACACAGAGTAGGC, respectively, and the *Cx3cr1* forward and reverse primers were TCGTCTTCACGTTCGGTCTG and CTCAAGGCCAGGTTCAGGAG, respectively. PCR amplification was performed using a 50 μL reaction mix containing 25 µL of GoTaq 2x MasterMix (Promega; Madison, WI, Cat# M7132), 10 μM each of the corresponding forward and reverse primers, 4 μL of cDNA template, and nuclease-free water added to 50 μL. For both target transcripts, the PCR amplification consisted of a two-minute initial denaturation at 95°C, 30 cycles of 30 second denaturation at 95°C, one-minute annealing at 55°C, one-minute extension at 72°C, and a final five-minute extension at 72°C. PCR products were then prepared with 2 μL of 6x TriTrack DNA loading buffer (Thermo Fisher Scientific, Cat# R1161) and 10 μL sample and run in a 2% agarose Tris-borate-EDTA (TBE) gel. Product size was visualized by emission at 600 nM in a LI-COR Oddysey Fc imager (LI-COR Biotechnology; Lincoln, NE) and evaluated by comparison to a GeneRuler 100 bp Plus DNA ladder (Thermo Fisher Scientific, Cat# SM0321).

### Phase-contrast Imaging

Phase-contrast images of cells from three randomly chosen fields per treatment were acquired just prior to extracellular flux measurements using an EVOS® FL Auto Cell Imaging System (Thermo Fisher Scientific; Waltham, MA). Manual cell counting was performed using the Fiji software (National Institutes of Health, Bethesda, MD) cell counting tool. Cell counts per field ranged from 25 to 281 cells and the cell number per well was estimated based on the field area (0.00142163 cm^2^) relative to the total well area (0.275 cm^2^). The mean of three fields was used as the cell count for each treatment group.

### Seahorse XF24 Extracellular Flux Analysis

Two experimental paradigms were used. In the first, HAPI microglial cells were non-activated or stimulated for 18 hours with LPS (100 ng/mL), interferon-gamma (IFN-γ; 10 ng/mL), Universal™ Type I interferon (hereafter, referred to as IFN-α; 10 ng/mL), or LPS in combination with IFN-γ or IFN-α at the same concentrations. In the second, non-activated or LPS/IFN-γ-treated HAPI cells were incubated for 18 hours in the absence or presence of N-3-(aminomethyl)benzylacetamidine (1400W, 50 µM), an inhibitor of iNOS (Garvey et al., 1997). The vehicle for these treatments was water and the volumes added were trivial compared to the volume of aqueous medium. The HAPI oxygen consumption rates (OCR) and extracellular acidification rates (ECAR) were assessed using a Seahorse XF24 Extracellular Flux Analyzer (Agilent Technologies) as previously described (Gerencser et al., 2009; Jaber et al., 2018). Measurements were performed in a pH 7.4 artificial cerebrospinal fluid assay medium comprised of 120 mM NaCl, 3.5 mM KCl, 1.3 mM CaCl2, 0.4 mM KH2PO4, 1 mM MgCl2, 5 mM 4-(2-hydroxyethyl)-1-piperazineethanesulfonic acid (HEPES), 15 mM glucose, and 4 mg/mL fatty acid-free bovine serum albumin.

Three baseline OCR and ECAR were acquired, and then the following reagents at the indicated final concentrations were added sequentially via injection ports A through C, with two rate measurements after each injection: Port A – carbonyl cyanide 4-(trifluoromethoxy)phenylhydrazone (FCCP, 4 µM) and pyruvate (10 mM), Port B – 2-(4-Carboxyphenyl)-4,4,5,5-tetramethylimidazoline-1-oxyl-3-oxide sodium salt (cPTIO, 200 µM), and Port C – antimycin A (1 µM). The sodium pyruvate (Sigma-Aldrich Cat# P2256) was prepared fresh from powder immediately prior to each experiment. OCR and ECAR were normalized to the cell count for each treatment group. Measurement cycles consisted of 2-minute (min) mix, 1.5- or 2-min wait, and 2-min measure periods. In two paradigm 1 experiments, two additional 2-min mix and 2-min wait periods were mistakenly introduced after the first two basal rate measurements, leading to a difference in the overall time course even though the timing of measurements 3-9 encompassing all drug injections was identical in the five experiments. Therefore, averaged OCR data are plotted against measurement number rather than time. For these paradigm 1 experiments, basal mitochondrial OCR was calculated by subtracting the antimycin A-insensitive OCR (measurement number 8) from the third baseline OCR measurement, maximal mitochondrial OCR was calculated by subtracting the antimycin A-insensitive OCR from the FCCP plus pyruvate-induced OCR (measurement number 4), and the third baseline ECAR measurement was considered to represent the basal ECAR.

### Nitric Oxide (NO) Quantification

NO release was evaluated using two experimental paradigms. The first was identical to the first Seahorse extracellular flux experiment paradigm, and NO release was quantified from the cell culture media of the Seahorse plate prior to the bioenergetics measurements. In the second, HAPI cells plated overnight at 50,000 cells per well in standard 24-well plates were non-activated or stimulated for 18 hours with LPS (100 ng/mL) plus or minus various concentrations of recombinant rat or mouse IFN-γ (2.5-10 ng/mL). To evaluate the amount of NO release from cells in each experimental paradigm, nitrite, a stable, non-volatile breakdown product of NO, was quantified in cell culture media by Griess assay (Thermo Fisher Scientific) according to the manufacturer’s protocol.

### Statistics

A two-way analysis of variance (ANOVA) with treatment and experiment number as factors was used to test for a significant effect of treatment on OCR, ECAR, or cell number (all n=5) using GraphPad Prism software version 10.4.1 (GraphPad, Boston, MA). Tukey’s post hoc analysis was used for pairwise comparisons among multiple groups. A two-way ANOVA with repeated measures and Šídák’s multiple comparisons test was used to test for a significant effect of cPTIO on OCR by comparing the OCR immediately before and after cPTIO injection. Linear regression analysis was used to test for a correlation between cPTIO-induced OCR and cell number. For all analyses, a p-value of <0.05 was considered significant.

## Results

### HAPI cells are microglia-like mouse cells

We obtained HAPI cells from the original source reporting a primary rat cell culture origin (Cheepsunthorn et al., 2001). Although HAPI cells are now identified as mouse cells in the Register of Misidentified Cell Lines, their true origin is unknown. To authenticate HAPI cells as mouse cells and identify if they have a match in the ATCC cell database, we submitted HAPI cells to ATCC’s Mouse Cell Line Authentication Service. This service is based on STR profiling using 18 STRs by a multiplex PCR assay established by the Mouse Cell Line Authentication Consortium (Almeida et al., 2019). STRs are short sequences of DNA that are repeated in tandem at specific locations (loci) in the genome. The number of repeats varies greatly among cell lines, making STRs highly useful for identification by providing a cell line “fingerprint.” STR profiling revealed that HAPI cells are a 100% STR match for SIM-A9 cells, a mouse microglial cell line in the ATCC database that was reported to originate from a postnatal day 1 primary mouse microglial culture through spontaneous immortalization (Nagamoto-Combs et al., 2014) (Table 1; Figure S1). Neither of two human STR markers were detected.

**Table 1.** Comparative short tandem repeat (STR) allele results for submitted HAPI cells and the SIM-A9 cell record present in the ATCC Mouse STR Database.

| Locus | HAPI Results | SIM-A9 Reference Profile |
| --- | --- | --- |
| 18-3 | 16,17 | 16,17 |
| 4-2 | 20.3 | 20.3 |
| 6-7 | 15 | 15 |
| 19-2 | 13 | 13 |
| 1-2 | 19 | 19 |
| 7-1 | 26.2 | 26.2 |
| 1-1 | 16,17 | 16,17 |
| 3-2 | 14 | 14 |
| 8-1 | 16 | 16 |
| 2-1 | 16 | 16 |
| 15-3 | 22.3,23.3 | 22.3,23.3 |
| 6-4 | 18 | 18 |
| 11-2 | 16 | 16 |
| 17-2 | 15 | 15 |
| 12-1 | 17 | 17 |
| 5-5 | 17 | 17 |
| X-1 | 27 | 27 |
| 13-1 | 17 | 17 |

The microglia-like HAPI cell identity was originally established through multiple microglial cell markers that are also expressed by macrophages, e.g., CD11b and GLUT5 (Cheepsunthorn et al., 2001). Subsequent to the HAPI cell line’s initial description, the microglia-selective marker TMEM119 was identified and it has been widely used to distinguish between microglia and macrophages (Bennett et al., 2016; Butovsky et al., 2014; Haage et al., 2019; Ye et al., 2023; Zrzavy et al., 2018). Although not entirely selective for microglia *in vivo* (Grassivaro et al., 2020; Vankriekelsvenne et al., 2022), its expression can distinguish between microglia and macrophages *in vitro* (Bennett et al., 2016; Butovsky et al., 2014). To further support that HAPI cells have a microglia-like cell identity, we conducted immunocytochemistry for TMEM119 on fixed HAPI cells. Universal staining of the HAPI cell population was observed (Figure 1A), supporting that HAPI cells have a microglial origin.

**Figure 1.**
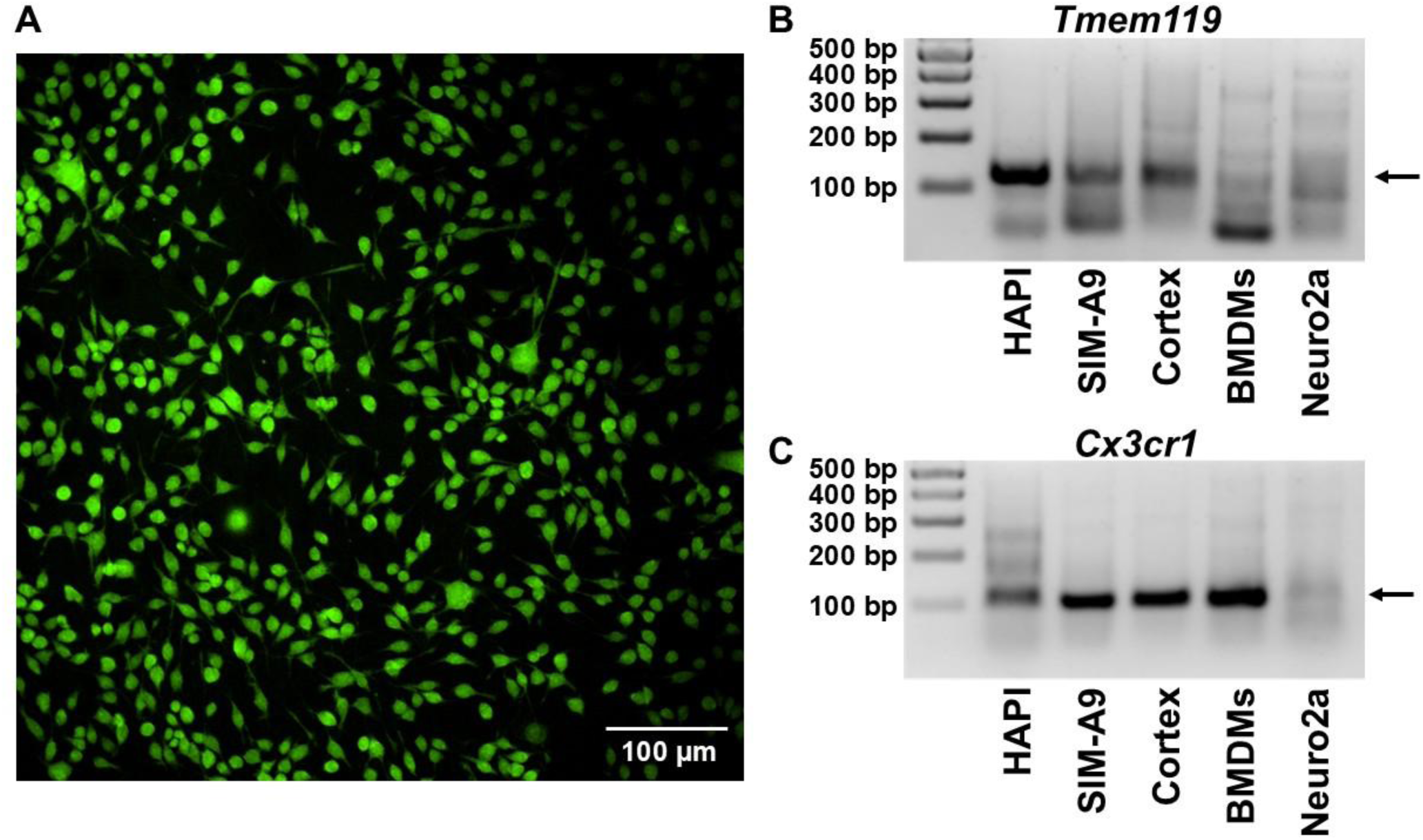
HAPI cells express microglia-selective markers. (A) Representative image of HAPI cells immunostained for TMEM119. (B) Detection of *Tmem119* polymerase chain reaction (PCR) product in sample prepared from HAPI cells, SIM-A9 cells, mouse cortex tissue, mouse bone marrow-derived macrophages (BMDMs), or Neuro2a cells. An arrow designates the band corresponding to the predicted product size of 119 base pairs (bp). (C) Detection of *Cx3cr1* PCR product in the same samples evaluated in B. An arrow designates the band corresponding to the predicted 109 bp product.

Because we were unable to identify any TMEM119 antibodies that are knockout-validated, we also investigated *Tmem119* expression at the mRNA level, given that the necessary homology of primers supports the specificity of amplified products. PCR amplification of complementary DNA using primers specific for microglia-selective *Tmem119* or for microglia- and macrophage-expressed *Cx3cr1* was carried out on samples isolated from HAPI cells, SIM-A9 mouse cells, mouse cortex (a microglia-containing positive control), primary mouse bone marrow-derived macrophages, and Neuro2a mouse neuroblastoma cells (Graham et al., 1974) (a negative control). We found that HAPI cells and SIM-A9 cells, but not BMDMs or Neuro2a cells, exhibited a *Tmem119* PCR product of the correct size, which was confirmed using microglia-containing mouse cortex as a positive control (Figure 1B). In contrast, the microglia-/macrophage-selective marker *Cx3cr1* was expressed by BMDMs as well as HAPI and SIM-A9 cells, whereas negligible PCR product was amplified from Neuro2a cells (Figure 1C). HAPI cells exhibited morphological changes in response to classical pro-inflammatory stimuli (Figure 2), including the formation of thin protrusions with phagocytic cups (arrowheads) that were most pronounced with IFN-γ treatment. Collectively, these results support that HAPI cells are a SIM-A9-related mouse microglia-like cell line.

**Figure 2.**
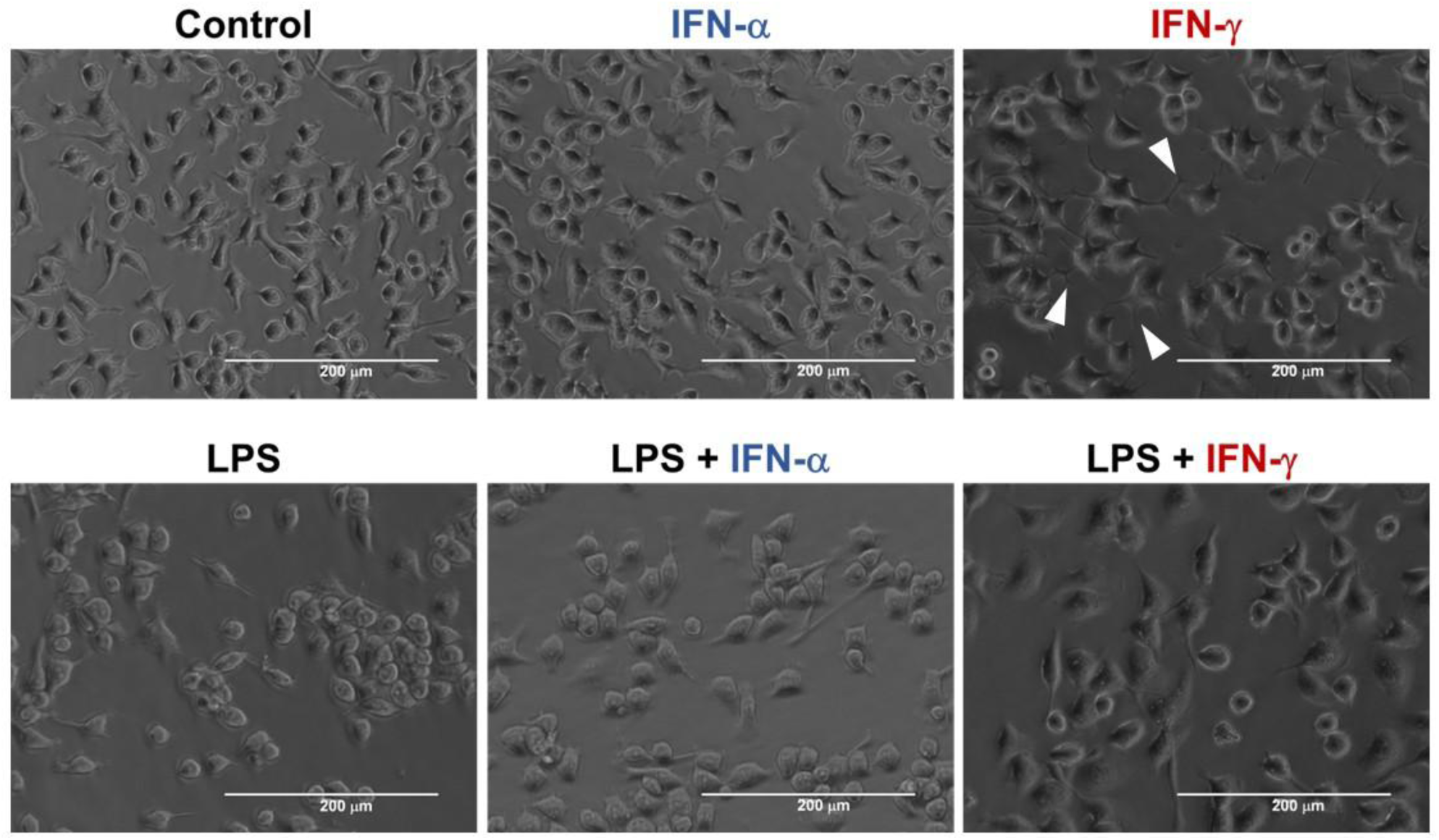
HAPI cells undergo morphology changes in response to various pro-inflammatory stimuli. Representative phase-contrast images of unstimulated HAPI cells (control) and HAPI cells following 18 hours of stimulation with the toll-like receptor 4 agonist lipopolysaccharide (LPS, 100 ng/mL), the type I interferon, interferon-alpha (IFN-α, 10 ng/mL), the type II interferon, interferon-gamma (IFN-γ, 10 ng/mL), or combined LPS/IFN-α or LPS/IFN-γ at the same concentrations. The white arrowheads denote examples of protrusions with phagocytic cups.

### LPS-, IFN-α-, LPS plus IFN-γ-, or LPS plus IFN-α-stimulation suppresses mitochondrial bioenergetic function in HAPI cells

Primary microglia (Jaber et al., 2018; Orihuela et al., 2016) and macrophages (Bailey et al., 2019; Palmieri et al., 2020; Van den Bossche et al., 2016) undergo a shift in energy metabolism from oxidative phosphorylation to glycolysis upon pro-inflammatory stimulation with LPS plus IFN-γ (LPS/IFN-γ) or LPS alone. Immortalized BV-2 microglial cells exhibit immunometabolism that is similar to primary microglia (Chenais et al., 2002; Moss and Bates, 2001; Orihuela et al., 2016; Voloboueva et al., 2013), but the pro-inflammatory bioenergetic responses of HAPI cells have yet to be characterized.

We employed Seahorse extracellular flux analysis to measure cellular OCR and ECAR — indicators of oxidative phosphorylation and glycolysis, respectively — after 18 hours of treatment with LPS, IFN-γ, IFN-α, LPS/IFN-γ, or LPS plus IFN-α (LPS/IFN-α). Following the acquisition of basal OCR and ECAR, a combination of the mitochondrial uncoupler FCCP and the cell permeable Complex I substrate pyruvate was added to induce maximal OCR, the NO scavenger cPTIO was injected to test for reversibility of respiratory inhibition, and the electron transport chain Complex III inhibitor antimycin A was added to enable subtraction of non-mitochondrial OCR. Rat IFN-γ was used for these bioenergetics experiments based on the initial presumption that HAPI cells are rat cells. However, a titration with rat and mouse IFN-γ revealed that the employed rat IFN-γ concentration (10 ng/mL) was sufficient to induce a maximal NO response equivalent to that induced by the mouse cytokine (Figure S2A). The high proliferation rate of HAPI cells made it difficult to achieve a consistent cell density at the time of assay since small inconsistencies in plating density were amplified over the experimental time-course (Figure 3A).

**Figure 3.**
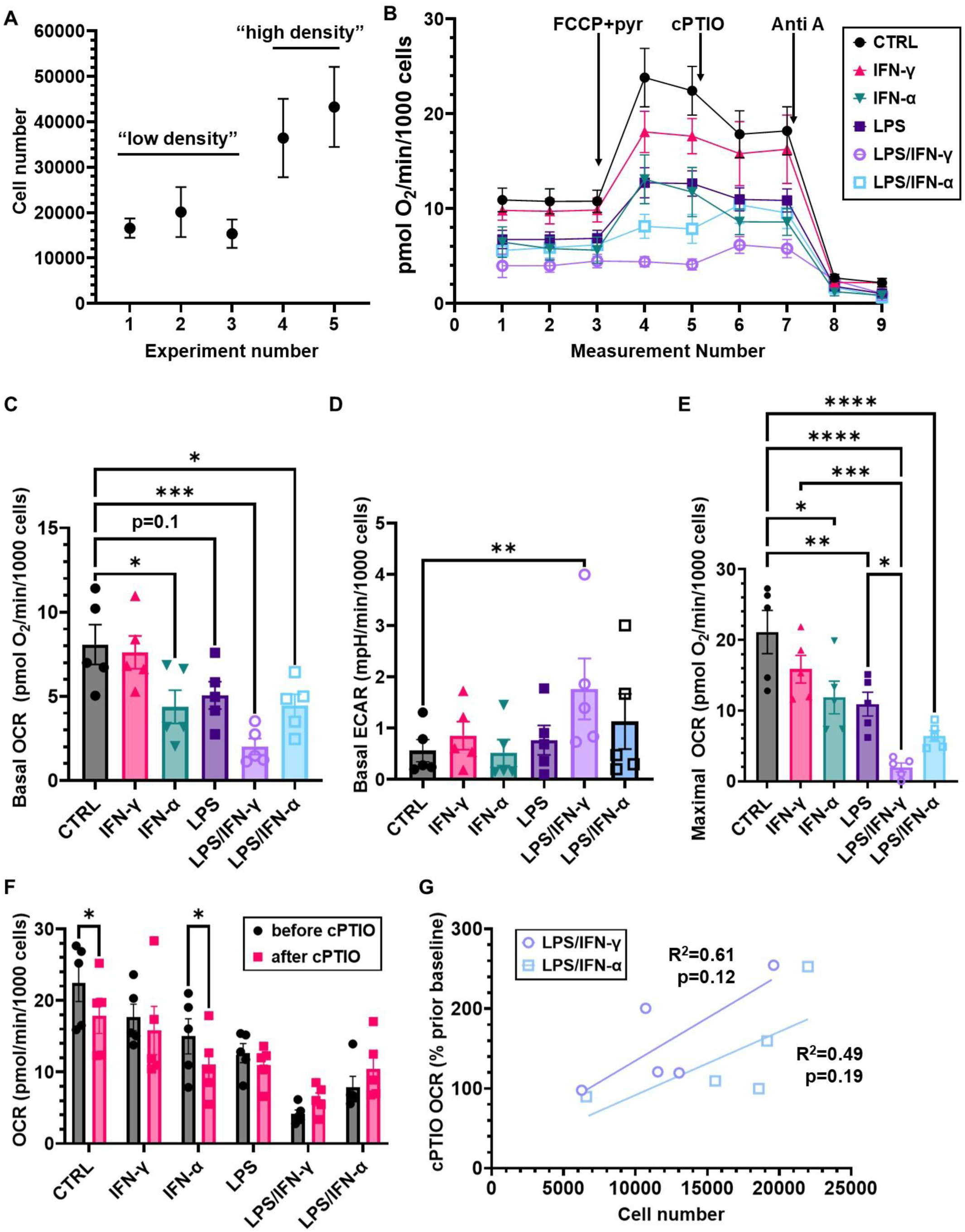
HAPI cellular bioenergetic function is altered by various pro-inflammatory stimuli. (A) Cell number of the non-activated control (CTRL) group plotted across the five experiments, showing the variability binned into low- and high-cell-density categories. (B) Average HAPI cell oxygen consumption rate (OCR) measurements normalized to cell number following 18 hours of stimulation with lipopolysaccharide (LPS, 100 ng/mL), interferon-gamma (IFN-γ, 10 ng/mL), interferon-alpha (IFN-α, 10 ng/mL), or combined LPS/IFN-γ or LPS/IFN-α at the same concentrations. The results are mean ± SEM of n=5 biological replicates, with each experiment consisting of 2-3 technical replicates. The serial additions of the uncoupler FCCP (4 µM) + pyruvate (pyr, 10 mM), the nitric oxide scavenger cPTIO (200 μM), and the Complex III inhibitor antimycin A (Anti A, 1 μM) are indicated by arrows. (C) Basal mitochondrial OCR normalized to cell number calculated from the data shown in B. (D) Basal extracellular acidification rate (ECAR) normalized to cell number. ECARs were acquired simultaneously to OCR for the experiment shown in B. (E) Maximal mitochondrial OCR normalized to cell number calculated from the data shown in B. The data in panels C-E were analyzed by two-way analysis of variance (ANOVA) with treatment and experiment number as factors, followed by Tukey’s post hoc analysis. Experiment number had a significant effect on basal OCR (p<0.05) and basal ECAR (p<0.0001), while only a strong trend was observed on maximal OCR (p=0.06). (F) OCRs before and after cPTIO addition for the indicated treatment groups for the experiment shown in B. The data in panel F were analyzed by two-way ANOVA with repeated measures and Šídák’s multiple comparison’s test. (G) Linear regression analysis of cPTIO-induced OCR versus cell number for the LPS/IFN-γ and LPS/IFN-α treatment groups, with R^2^ and p-values indicated on the graph. The p-values indicate the probability of a non-zero slope of the linear fits. *p<0.05, **p<0.01, ***p<0.001, ****p<0.0001.

To account for the potential influence of this and other sources of variability, the individual experiment number was included as a factor when using ANOVA to test for a main effect of treatment on bioenergetics parameters. The mean OCR traces are shown in Figure 3B and examples of low-and high-cell-density experiments (experiment number 2 and 4, respectively) are shown in Figure S3.

The basal mitochondrial OCR was significantly decreased by LPS when added in combination with either IFN-γ or IFN-α and, also, decreased by IFN-α treatment alone (Figure 3C). Treatment with LPS alone exhibited a trend toward basal OCR inhibition (p=0.1), while the basal OCR upon IFN-γ stimulation was unchanged from the non-activated control. The basal ECAR was significantly increased only by combined LPS/IFN-γ treatment (Figure 3D). However, the variable cell density may have obscured IFN-α-, LPS-, and LPS/IFN-α-induced ECAR increases, as trends were apparent when the low- and high-density experiments were plotted separately (Figure S4).

All pro-inflammatory stimuli except for IFN-γ significantly impaired maximal OCR (Figure 3E), which was measured following the addition of FCCP with excess substrate. Although IFN-γ added individually did not affect OCR, it synergistically increased the LPS-induced maximal OCR suppression, consistent with its ability to synergistically enhance LPS-induced NO production (Papageorgiou et al., 2016) (Figure S2B). In contrast, IFN-α alone caused maximal OCR suppression and the level of reduction seen with combined LPS/IFN-α treatment did not reach significance compared with LPS or IFN-α treatment alone despite its similar ability to synergistically increase NO (Figure S2B). LPS/IFN-γ treatment caused a significant reduction in cell number compared to the non-activated control group, consistent with its reported ability to induce primary microglia cell death (Mayo and Stein, 2007), and LPS/IFN-α treatment also led to a significantly reduced cell count (Figure 4).

**Figure 4.**
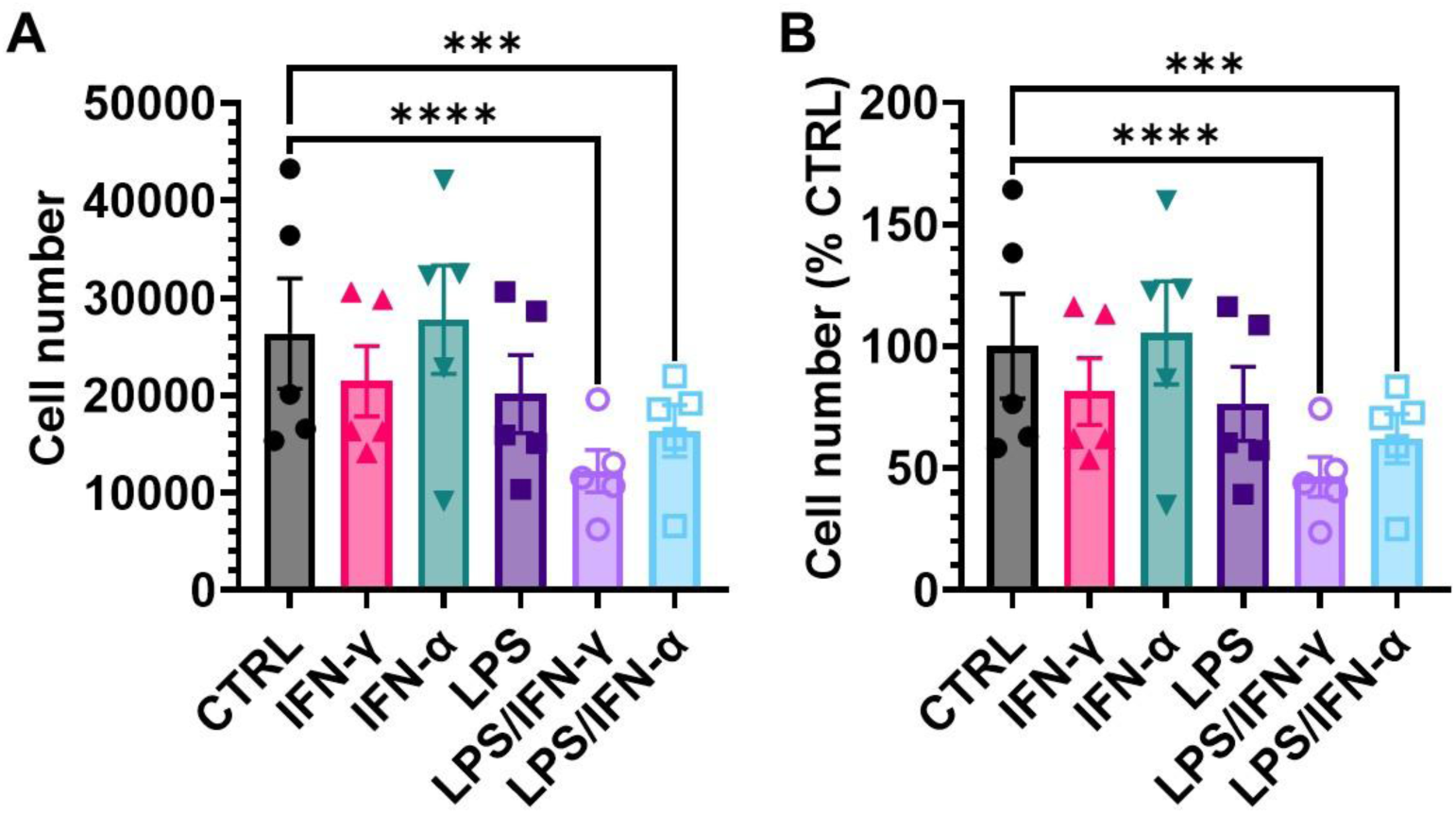
Eighteen-hour activation of HAPI cells with lipopolysaccharide plus interferon-gamma (LPS/IFN-γ) or LPS plus interferon-alpha (LPS/IFN-α) reduces cell number compared to the non-activated control (CTRL) group. HAPI cells were treated with the following stimuli prior to the determination of cell number by manual cell counting of three representative fields per treatment: LPS (100 ng/mL), IFN-α (10 ng/mL), IFN-γ (10 ng/mL), LPS/IFN-γ (100 ng/mL / 10 ng/mL, respectively), or LPS/IFN-α (100 ng/mL / 10 ng/mL, respectively). Cell number is shown in (A) while the results in (B) express the cell number as a percentage of the CTRL group mean. The results are mean ± SEM of n=5 biological replicates. The data were analyzed by two-way analysis of variance with treatment and experiment number as factors, followed by Tukey’s post hoc analysis. Both the treatment and the experiment number had a significant effect (p<0.0001 each), and there was a significant interaction between the two (p<0.05). ***p<0.001, ****p<0.0001.

Inhibiting or ablating the NO-producing enzyme iNOS reduces the level of mitochondrial respiratory suppression by LPS ± IFN-γ in macrophages or microglia (Bailey et al., 2019; Orihuela et al., 2016; Palmieri et al., 2020; Van den Bossche et al., 2016). To evaluate whether any of the respiratory suppression caused by IFN-α, LPS, LPS/IFN-γ, or LPS/IFN-α was acutely reversible by removing NO, we introduced the NO scavenger cPTIO following the FCCP-triggered induction of maximal OCR. CPTIO did not significantly rescue maximal OCR impaired by any of the pro-inflammatory treatments across the five experiments, although it appeared to stimulate OCR in LPS/IFN-γ- or LPS/IFN-α-treated cells in a subset of the experiments (e.g., the LPS/IFN-α-treated cells in Figure S3B). Quantification of NO production from the same cells used for the Figure S3B OCR measurements revealed that both IFN-γ and IFN-α synergistically enhanced NO production upon LPS treatment, but to a significantly greater extent with IFN-γ even though the rescue trend was seen for the LPS/IFN-α treatment (Figure S2B). Linear regression analysis was used to test for a relationship between cPTIO-induced OCR and cell density for the LPS/IFN-γ and LPS/IFN-α treatments (Figure 3G). Although the greatest cPTIO OCR stimulation was observed at the highest cell density for each treatment, there was not a significant cell number-OCR correlation for either stimulus. Despite cPTIO’s failure to rescue maximal OCR, the iNOS inhibitor 1400W partially prevented the HAPI cell LPS/IFN-γ-induced maximal OCR impairment (Figure 5), consistent with the partial iNOS-dependence of respiratory suppression observed in primary microglia and macrophages. It was confirmed that 1400W completely prevented NO production by LPS/IFN-γ-stimulated cells and did not affect the OCR of non-activated control cells (data not shown). Collectively, these bioenergetics experiments indicate that the pro-inflammatory immunometabolism of HAPI cells recapitulates that of primary microglia, as well as macrophages, supporting the continued use of HAPI cells as a microglial model for mechanistic studies.

**Figure 5.**
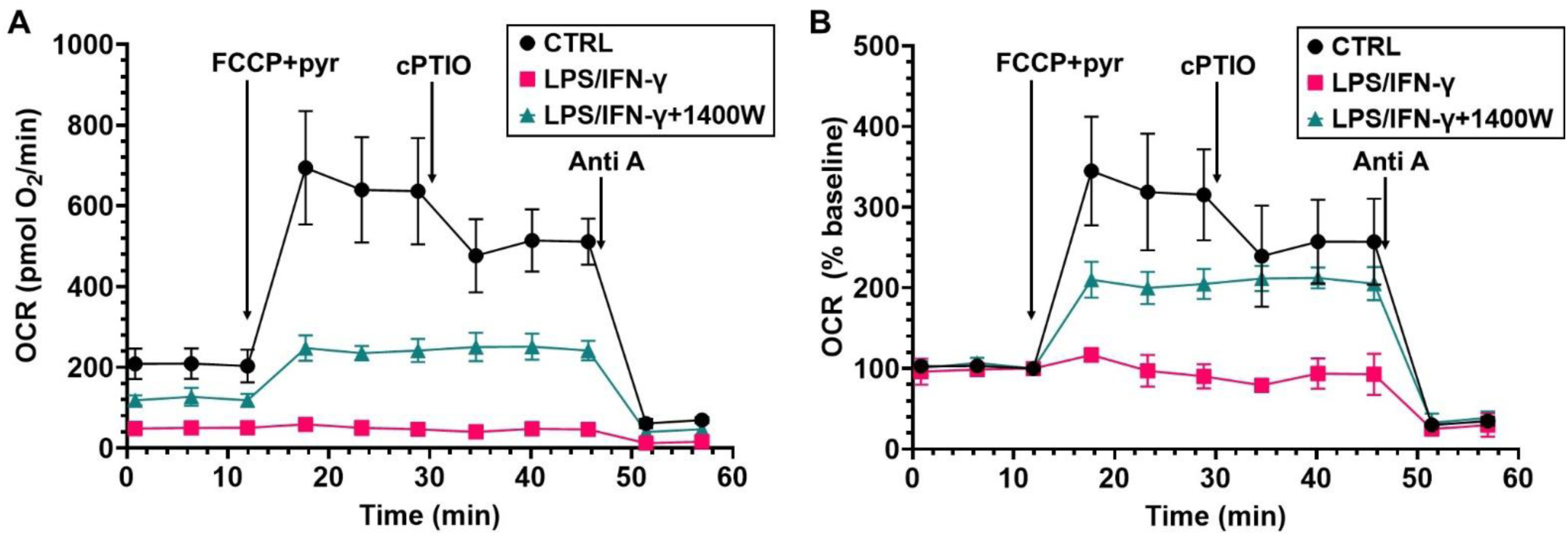
Respiratory suppression induced by combined lipopolysaccharide/interferon-gamma (LPS/IFN-γ) stimulation is partly attenuated by an inhibitor of inducible nitric oxide synthase (iNOS). (A) HAPI cell oxygen consumption rate (OCR) measurements from an unstimulated control (CTRL) group or following 18 hours of stimulation with LPS/IFN-γ (100 ng/mL / 10 ng/mL, respectively) in the absence or presence of the iNOS inhibitor 1400W (50 µM). The serial additions of FCCP (4 µM) + pyruvate (pyr, 10 mM), cPTIO (200 μM), and antimycin A (Anti A, 1 μM) are indicated by arrows. (B) The data in A expressed as a percentage of the baseline OCR (third measurement) because cell counts were not done for this experiment. Baseline normalization removes the influence of cell number on OCR, illustrating that the recovery of respiratory capacity is incomplete. The results are mean ± SD of n=3 technical replicates and are representative of n=7 similar experiments done in triplicate. Because this experiment was done as part of another study with additional comparisons, only the representative trace is shown, and the results will be reported in more detail elsewhere.

## Discussion

The usage of HAPI cells to model pro-inflammatory microglial responses *in vitro* has been widespread since their initial description as spontaneously immortalized rat microglia in 2001 (Cheepsunthorn et al., 2001). However, their rat microglia-like identity was called into question in 2021 when they were listed in the Register of Misidentified Cell Lines database as mouse cells of unknown origin. Using the gold-standard cell line authentication method of STR profiling based on 18 different short tandem repeats (Almeida et al., 2019), we discovered that HAPI cells are an exact match to SIM-A9 cells, a mouse cell line also reported to arise via spontaneous immortalization from a primary microglia culture (Nagamoto-Combs et al., 2014). To further support the mouse microglia-like identity of the HAPI cell line, we demonstrated mRNA and protein expression of the microglia-selective marker TMEM119 and further characterized their morphological and bioenergetic responses to LPS and IFN-γ, both individually and in combination. We also investigated the bioenergetic responses of a microglial cell line to IFN-α for the first time.

STR profiling has proven to be a highly accurate method of cell line authentication. An earlier STR profiling method based on only nine STRs was able to distinguish among three different Balb/c-derived cell lines (Almeida et al., 2014), indicating its high discriminatory potential. Therefore, the 100% eighteen-STR match strongly suggests that HAPI cells and SIM-A9 cells—which are not listed in the Register of Misidentified Cell Lines database—have a common origin. It is unlikely that HAPI cells are simply mischaracterized SIM-A9 cells given that HAPI cells appeared in the literature 13 years prior to when SIM-A9 cells were reported. However, although SIM-A9 cells are not listed in the Register of Misidentified Cell Lines database, given the chronology, it is possible that SIM-A9 cells are mischaracterized HAPI cells, and HAPI cells are mischaracterized from another microglial cell line. An alternative possibility is that both HAPI cells and SIM-A9 cells are descended from the same immortalized microglial cell line. Regardless, the expression of the microglia-selective marker TMEM119 by both HAPI cells and SIM-A9 cells, combined with that of *Cx3cr1* and previously described markers (Cheepsunthorn et al., 2001; Nagamoto-Combs et al., 2014), strongly support a microglial cell origin for both cell lines.

Another goal of this study was to provide an initial characterization of HAPI bioenergetic responses to pro-inflammatory stimuli, supporting the use of HAPI cells for such investigations. Given the involvement of TLR4-dependent signaling in acute brain injury and multiple neurodegenerative diseases, as well as the presence of IFN-γ-producing leukocytes in many of the same conditions, we studied the responses of HAPI cells to the TLR4 agonist LPS and the Type II interferon, IFN-γ. First, we reproduced previous findings that HAPI cells, like primary rodent microglia, respond to LPS/IFN-γ treatment with NO production and a shift toward ameboid-like morphology (Cheepsunthorn et al., 2001; Chuang et al., 2013; Dallas et al., 2013; Hara et al., 2014; Hara et al., 2024; Hara et al., 2011; Horvath et al., 2008; Jantaratnotai et al., 2022; Jantaratnotai et al., 2006; Jantaratnotai et al., 2013; J. Li et al., 2022; Lo et al., 2008; McFarland et al., 2018; Pawate et al., 2004; Sheng et al., 2011; Thampithak et al., 2009; Tocharus et al., 2012). Using the iNOS inhibitor 1400W, we then showed that HAPI cells, like primary rodent microglia/macrophages and BV-2 microglial cells, exhibit a partially NO-dependent suppression of mitochondrial oxygen consumption after LPS and IFN-γ treatment (Bailey et al., 2019; Orihuela et al., 2016; Palmieri et al., 2020; Van den Bossche et al., 2016).

Aside from IFN-γ’s role in neurodegenerative disease, Type I interferon signaling has recently been implicated in diverse neurological conditions, including TBI (Barrett et al., 2020; Todd et al., 2023), Alzheimer’s disease (Roy et al., 2022; Roy et al., 2020), neuropsychiatric lupus (Aw et al., 2023; Zervides et al., 2025), and neuroinflammation induced by human immunodeficiency virus (Koneru et al., 2018). Characterization of the microglial responses to Type I interferons has included transcriptional changes, morphological alterations, phagocytic activity, and inflammatory phenotypes (Aw et al., 2020; W. Li et al., 2018; McDonough et al., 2017), but we were unable to find any studies examining mitochondrial parameters. Therefore, we extended HAPI cell immunometabolism characterization beyond the well-studied LPS/IFN-γ paradigm to investigate the effects of a Type I interferon. We discovered that IFN-α enhances LPS-induced NO release, reduces cell number in combination with LPS, and reduces basal and maximal mitochondrial OCR, either with or without concomitant LPS stimulation. A limitation of this study is that we did not also determine the response of primary microglia to IFN-α, so the extent to which these results recapitulate the behavior of primary cells is unknown. However, the Type I interferon, IFN-β, was reported to inhibit mitochondrial respiration in primary macrophages, both individually and in combination with LPS (Olson et al., 2021), and IFN-α was reported to inhibit the respiration of a B lymphoblast cell line (Lewis et al., 1996; Lou et al., 1994), demonstrating that Type I interferons can suppress mitochondrial bioenergetic function in multiple immune cell types.

NO can inhibit the mitochondrial electron transport chain through multiple mechanisms that include S-nitrosylation (Borutaite et al., 2000; Clementi et al., 1998), reversible competition with oxygen at Complex IV (Brown et al., 1995; Brown and Cooper, 1994; Cleeter et al., 1994; Schweizer and Richter, 1994), and irreversible damage to respiratory chain complexes mediated by peroxynitrite, which is formed by the NO-superoxide reaction (Cassina and Radi, 1996; Lizasoain et al., 1996; Radi et al., 1994; Riobo et al., 2001). To evaluate whether any of the NO-mediated mitochondrial respiratory suppression by pro-inflammatory stimuli is acutely reversible, we tested whether the scavenger cPTIO acutely restores mitochondrial oxygen consumption. We failed to observe significant mitochondrial OCR rescue with cPTIO for any of the stimuli, although the NO scavenger appeared to show a very modest ability to reverse LPS/IFN-γ- or LPS/IFN-α-induced respiratory inhibition in a few individual experiments (e.g., see Fig. S3B). We hypothesized that inconsistent cell density may explain this variability, as the medium dissolved oxygen concentration is expected to be lower in higher-density cultures due to mitochondria-mediated oxygen depletion, which would allow NO to compete more effectively at Complex IV. We observed only weak trends toward a linear relationship between cell count and cPTIO-induced OCR for the LPS/IFN-γ or LPS/IFN-α treatment (p=0.12 or p=0.19, respectively). This does not rule out the possibility that a relationship was obscured by the potential inaccuracy of cell count estimates from randomly selected fields or that there is a cell density threshold for cPTIO responsiveness. However, because the possible cPTIO rescue was very modest even when observed and consistent cell densities were hard to achieve, we decided not to further investigate the possibility of NO-scavenging-mediated OCR rescue.

In summary, we showed that HAPI cells are a mouse microglial cell line that appears to have a common origin with the SIM-A9 mouse microglial line. Like BV-2 and primary rodent microglia, HAPI cells demonstrated suppression of mitochondrial oxygen consumption after LPS and IFN-γ treatment that was partly NO-mediated and not reversed by NO scavenging. The Type I interferon IFN-α also inhibited mitochondrial respiration in the absence and presence of LPS. Our microglia marker expression and immunometabolism studies support the continued use of HAPI cells to study rodent pro-inflammatory microglial signaling *in vitro*.

## Acknowledgements

The authors acknowledge support from the T32 Training Program in Integrative Membrane Biology, National Institutes of Health (NIH) grant number GM008181 (R.P.M.) and the University of Maryland School of Medicine Core Confocal Facility, which is supported by NIH grant number S10 OD026698.

## Data availability statement

The data supporting the findings of this study are available from the authors upon reasonable request.

