## Supplemental Material for "HAPI Cells are SIM-A9-related Mouse Microglial Cells Useful for *In Vitro* Modeling of Microglial Immunometabolism"

### Supplemental Figures

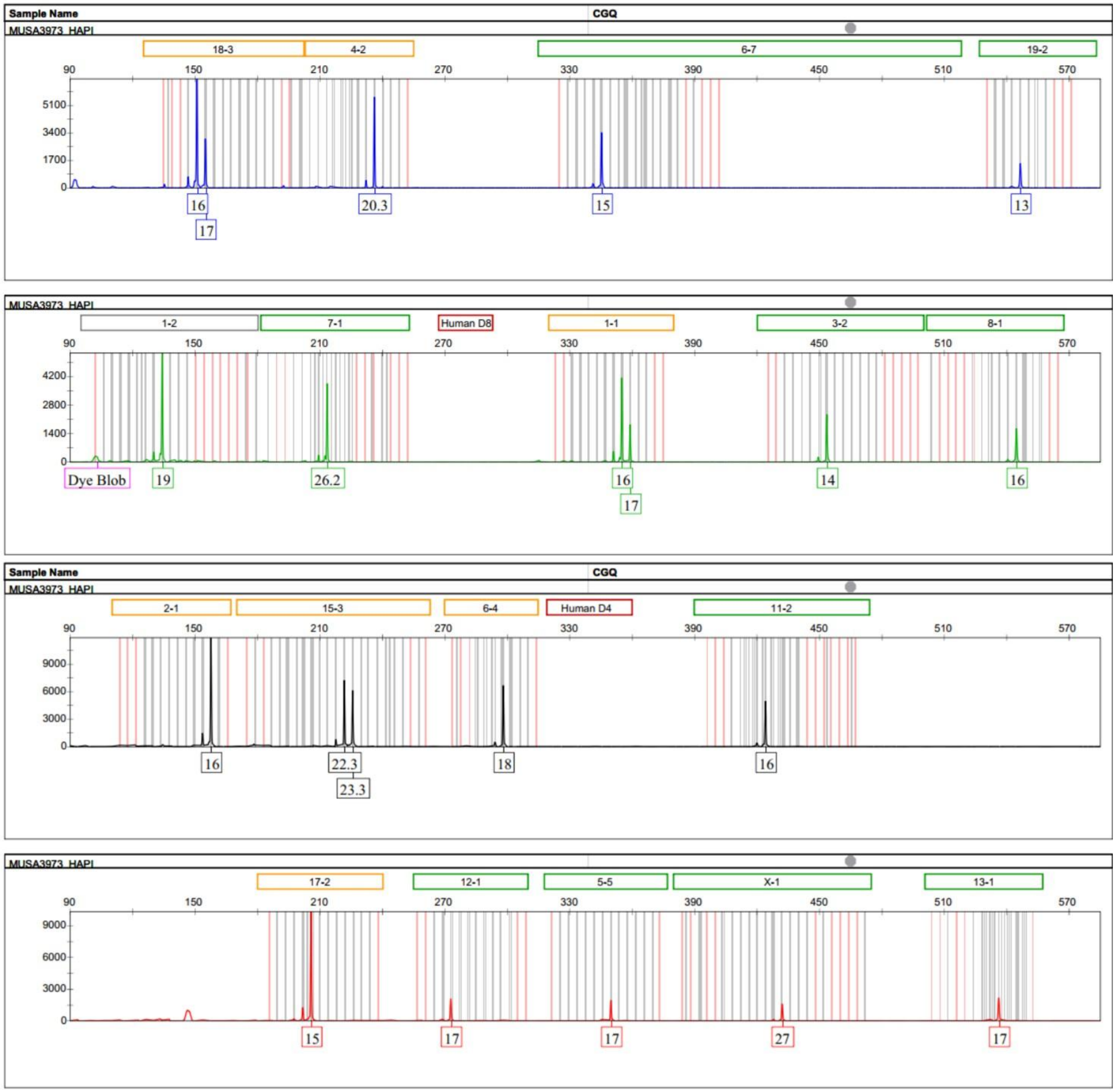


**Figure S1.** Electropherograms of each allele used for HAPI cell short tandem repeat (STR) profiling.

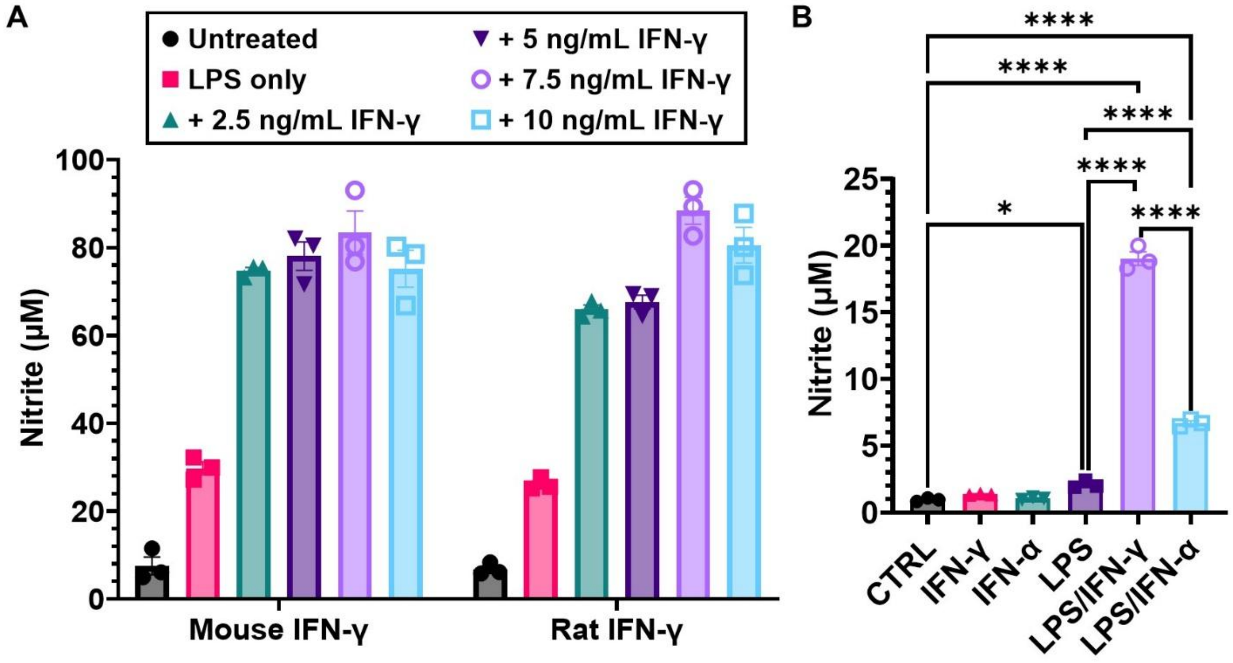


**Figure S2.** Either rat or mouse recombinant interferon-gamma (IFN-γ) or cross-species-activating recombinant interferon-alpha (IFN-α) synergistically enhance lipopolysaccharide (LPS)-induced nitric oxide (NO) release. NO release was measured by quantification of nitrite, the stable byproduct of NO, in cell culture media. (A) Titration of IFN-γ reveals that both mouse-origin and rat-origin IFN-γ induce maximal NO release when combined with LPS at the 10 ng/mL IFN-γ concentration used for bioenergetics experiments. LPS was present at 100 ng/mL in all but the untreated group. The results are mean ± SD of nitrite measurements done in triplicate. (B) Synergistic enhancement of LPS (100 ng/mL)-induced NO release by either IFN-γ (10 ng/mL) or IFN-α (10 ng/mL) in experiment number 4. The results are mean ± SD of nitrite measurements done in triplicate. The data were analyzed by one-way analysis of variance (ANOVA), followed by Tukey's post hoc test. *$p<0.05$, ****$p<0.0001$

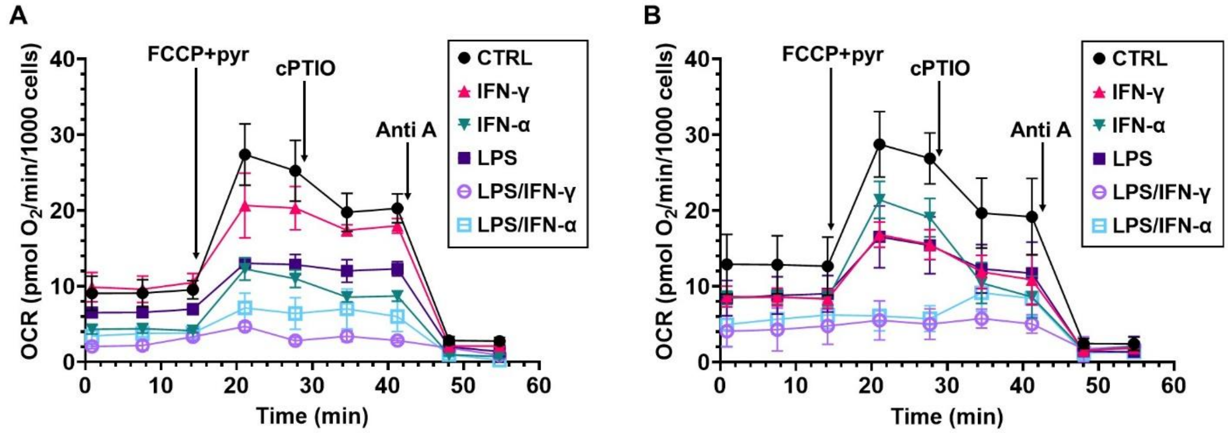


**Figure S3.** Example oxygen consumption rate (OCR) traces for low- and high-cell-density experiments following 18 hours of pro-inflammatory HAPI cell stimulation with individual and combined lipopolysaccharide (LPS) and interferon treatments. (A) Low cell density experiment number 2 results. (B) High cell density experiment number 4 results. In both A and B, OCR measurements were made following 18 hours of stimulation with LPS (100 ng/mL), interferon-gamma (IFN-γ, 10 ng/mL), interferon-alpha (IFN-α, 10 ng/mL), or combined LPS/IFN-γ or LPS/IFN-α at the same concentrations. The results are mean ± SD of measurements done in triplicate and are normalized to cell number. The serial additions of the uncoupler FCCP (4 μM) + pyruvate (pyr, 10 mM), the nitric oxide scavenger cPTIO (200 μM), and the Complex III inhibitor antimycin A (Anti A, 1 μM) are indicated by arrows.

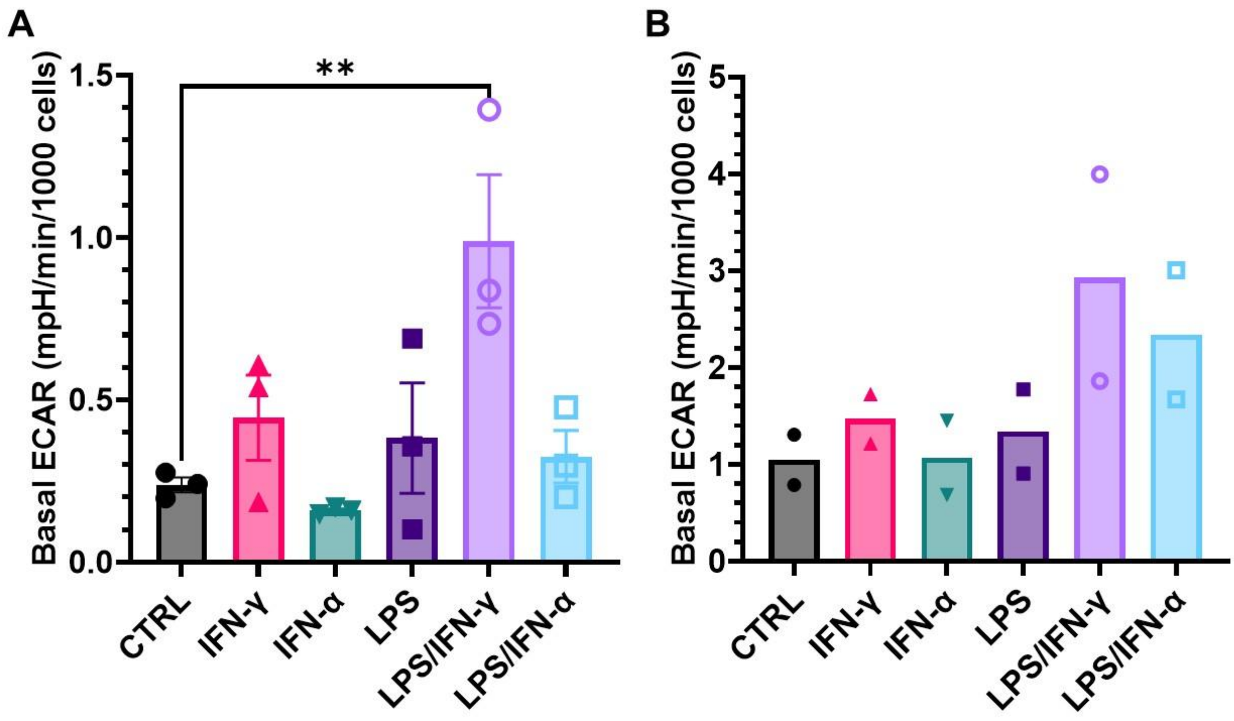


**Figure S4.** Basal extracellular acidification rate (ECAR) measurements for low- and high-cell-density experiments plotted separately reveal similar trends. ECAR measurements were made following 18 hours of stimulation with LPS (100 ng/mL), interferon-gamma (IFN-γ, 10 ng/mL), interferon-alpha (IFN-α, 10 ng/mL), or combined LPS/IFN-γ or LPS/IFN-α at the same concentrations. (A) Average basal ECAR normalized to cell number for the low cell density experiments (experiment numbers 1-3). The results are mean ± SD of measurements done in triplicate (B) Average basal ECAR normalized to cell number for the high cell density experiments (experiment numbers 4 and 5). The results in A are mean ± SEM of three biological replicates, with each treatment consisting of 2-3 technical replicates. The data were analyzed by one-way analysis of variance (ANOVA), followed by Tukey's post hoc test. The results in B are means of two biological replicates, with each treatment consisting of 3 technical replicates. $^{**}p<0.01$
